# Conjugation of bisphosphonates with scorpion venom enhanced apoptotic activities in pancreatic cancer cells

**DOI:** 10.1101/2022.08.30.505790

**Authors:** Nabil A. Alhakamy, Mohammed W. Al-Rabia, Waleed Y. Rizg, Adel F. Alghaith, Osama A. A. Ahmed, Usama A. Fahmy

## Abstract

**Background:** In recent times, prevalence and the incidence of pancreatic cancer (PC) have increased exponentially. Among of the major issues of PC are the asymptotic manifestation that delays diagnosis, and scarcity of therapeutic means.

**Methodology:** To provide an effective therapeutic approach, current study was executed, in which an optimized alendronate sodium (ALS) and scorpion venom (SV) nanoconjugate (ALS-SV nanoconjugate) was selected via experimental design applying size and zeta potential as the selected factors. Additionally, efficacy against PANC1 cells was determined by analysis of IC_50_, cell cycle, annexin V staining, Bcl-2, Bax, *p53*, etc.

**Results:** The particle size of the optimized nanoconjugate was found to be 28.4± 1.2 nm and the zeta potential to be +26.2± 0.8 mV. The cytotoxicity in terms of IC_50_ ALS-SV nanoconjugate was found better (2.73 ± 0.2 µg/mL) than ALS alone (22.3 ± 0.2 µg/mL) and SV alone (6.23 ± 0.4 µg/mL). Besides, outcomes of cell cycle analysis exhibited maximum efficacy for nanoconjugate in the G2-M phase. Annexin V staining, Bax, Bcl-2, *p53*, and caspase 3 estimation also showed superior apoptotic activity for ALS-SV nanoconjugate. However, as compared to ALS alone and SV alone, nanoconjugate showed elevated expression levels of TNF-α. The apoptotic activity was enhanced by the ALS-SV nanoconjugate as confirmed by MMP analysis.

**Conclusion:** Accordingly, the current study revealed an enhancement in the anti-PC effects of the ALS-SV nanoconjugate that is considered as a novel approach against PC.

## Introduction

Pancreatic cancer (PC) is among the major causes of death globally. As per the published report PC is the 14th common type of cancer and 7th cause of death [1,2]. Developed countries have a higher case of PC as compared to developing countries [1]. It was further reported that by the end of 2030, death due to PC might become the 2nd common cause. PC is associated with poor diagnosis and survival rate, as estimated by the various case studies and meta-analyses [3]. Among different types of PC, pancreatic adenocarcinoma alone is responsible for more than 90% of total PC [2,4]. In most cases (>70%), PA originates from the head of the pancreas, whereas >15% of PA originates from the body [5]. Moreover, PC is usually diagnosed at the advanced stage and becomes difficult to treat. A major challenge with the timely diagnosis of PC is multi-factorial in origin and non-specific signs and symptoms. As of now, resonant magnetic imaging (MRI), endoscopic ultrasound (EUS) CT scan and magnetic resonance cholangiopancreatography (MRCP) either alone or in combination are used for the diagnosis and prognosis of PC [6]. Currently, only serum cancer antigen 19-9 is an approved biomarker by the FDA for the management and treatment of PC, but this biomarker is associated with a lower predictive value [7]. Thus, mutation-based analysis was explored as an alternate biomarker, and KRAS mutation was a reliable and alternate one. Recently, volatile organic compounds (VOC) were reported to be a reliable biomarker for PC and offer a sensitivity of 84-100% [8]. As of now, only surgical intervention is an available therapeutic option to cure the PC. However, chemotherapeutic drugs are used as an adjuvant to prevent the reoccurrence and increase the overall survival rate [5].

Therefore, the lack of well-defined pharmacotherapeutics and lack of timely diagnostic tools creates serious problems for the management and treatment of PC. Anatomically, a major limitation in the treatment of PC is the deficit supply of blood vessels to the pancreas, and hence either radiotherapy or surgery is preferred [2]. Thus, targeting drug delivery appears to be a novel approach for the treatment of PC. A chemotherapeutic drug in a suitable nanocarrier is used to gain targeted therapeutic outcomes and minimal side effects compared to the free drugs. Furthermore, selective targeting to vascular endothelial growth factor (VEGF), RAS, or apoptotic signaling pathway in the acidic microenvironment in conjugated form is a novel approach [9].

Alendronate is one of the extensively used bisphosphonates in the management and treatment of osteoporosis. However, published reports have shown pro-apoptotic and anti-angiogenic effects against breast cancer and other types of cancer [10,11]. Additionally, bisphosphonates are already in use as an adjutant in cancer [12]. It is further important to highlight that among various drugs of class bisphosphonates, nitrogen-containing bisphosphonates have shown improved anticancer potential, and alendronate belongs to the nitrogen-containing bisphosphonates [10,11]. Furthermore, overexpressed cofilin is the hallmark of PC, and alendronate has been reported to downregulate the expression of cofilin in PC [13].

One of the novel approaches in the targeted drug delivery of pharmaceuticals in the case of PC is peptide-mediated drug targeting. Scorpion venom (SV) is an investigating natural product consisting of different types of disulfide-rich peptides (BMK-AGAP, BmTx3, AmmTx3, MgTX, etc.) that possess significant anticancer potential [14]. SV peptides have been previously reported to induce apoptosis, inhibit proliferation, angiogenesis, metastasis, and immunomodulatory effect in different types of cancers [14]. However, peptides suffer from the limitation of adequate stability, thermal degradation, fast metabolism, and a very short half-life [15]. Thus, we herein aim to deliver a combination of alendronate and SV to explore the anticancer potentiation of this combinatorial system in pancreatic cancer.

However, it is challenging to combine a pharmaceutical agent and peptide; hence, a nanoconjugagation approach was used. Nanoconjugate offers multiple advantages such as higher volume to surface area ratio, stability, biocompatibility, increased half-life, thermal stability, and reduced particle size that eventually helps in targeted drug delivery [11,16]. Additionally, peptides-drug conjugates have been reported with enhanced anticancer effects in various preclinical studies [16,17].

Therefore, the current study was designed to fabricate, characterize and optimize the Alendronate sodium (ALS)-SV nanoconjugates (ALS-SV nanoconjugates) using Box-Behnken response surface design. For this purpose, optimized nanoconjugate was optimized by analyzing dependent variables such as particle size (Y1) and zeta potential (Y2). Furthermore, the optimized nanoconjugate was studied for the anticancer effect in PANC1 cells, and various molecular parameters related to carcinogenesis were explored.

## Materials and Methods

### Materials

ALS, SV, colorimetric assay kit obtained from BioVision, Milpitas, CA, USA. In addition, a pancreatic cell line (PANC1) was procured from NCCS, Pune, India. All the other reagents and chemicals used were of the analytical grade.

### Experimental design for optimization of ALS-SV nanoconjugates

The Statgraphics plus, (Statgraphics Centurion XV version 15.2.05 software, Manugistic Inc., PA, USA) was used for designing Box-Behnken response surface and for formulating the ALS-SV nanoconjugate.

Independent variables’ statistical such as ALS: SV molar ratio (X1), sonication time (X3), and incubation time (X2) were recorded for the size (Y1), zeta potential (Y2) (Table 1).

**Table 1.** Factors’ levels and response constraints used in the Box-Behnken design for the optimization of ALS-SV nanoconjugates.

| Independent variables | Levels |  |  |
| --- | --- | --- | --- |
|  | -1 | 0 | +1 |
| X<sub>1</sub>: ALS: SV molar ratio | 1: 1 | 1: 8 | 1: 15 |
| X<sub>2</sub>: Incubation time (min) | 5 | 12.5 | 20 |
| X<sub>3</sub>: Sonication time (min) | 0.5 | 0.75 | 1.0 |
| Responses | Desirability constraints |  |  |
| Y1: Particle size (nm) | Minimize |  |  |
| Y2: Zeta potential (mV) | Maximize |  |  |

**Abbreviations:** ALS, alendronate sodium; SV, scorpion venom peptide

### Preparation of ALS-SV nanoconjugates

According to the runs obtained from the Box-Behnken-based Design Expert (Table 7), ALS-SV nanoconjugate of different concentrations of ALS: SV and with different incubation and sonication time were prepared. Briefly, ALS and SV were mixed in various proportions, and then it was dissolved in an aqueous phase (deionized water) in 20 mL. Next, the dissolved mixtures were incubated and sonicated for stated times according to runs obtained from the software (Table 1).

### Analysis of particle size and zeta potential

The size and zeta potential of the prepared ALS-SV nanoconjugates were determined by dissolving the formulation in double-distilled water (10-times dilution) using Zetasizer Nano ZSP (Malvern, UK) [11,18,19].

### Optimization of ALS-SV nanoconjugates

The effects of independent variables over dependent variables were examined, and an optimized nanoconjugate was selected after numerical analysis, which follows the desirability function approach. The aim of this optimization process was to obtain an ALS-SV nanoconjugate minimum particle size and maximum zeta potential. Next, the selected optimized nanoconjugate was prepared and further evaluated on several parameters such as cytotoxicity, cell cycle analysis, annexin V staining, caspase 3, etc.

### Optimized ALS-SV nanoconjugate cytotoxicity study via MTT assay

Optimized ALS-SV nanoconjugates was studied for cytotxic effects using the PANC1 cell line. PANC1 cells were overnight incubated in 96-well plates (5×103/cells/well). The plates were then exposed to ALS alone, SV alone, and nanoconjugate of ALS-SV and incubated for 48-h. The cells were then exposed to 10 µL of MTT solution (5.0 mg/ml) for 4 h at 37 °C. After that, the supernatants were collected and treated with 100 mL DMSO. The samples were detected at 570 nm using microplate readers [20].

### Cell cycle analysis of optimized ALS-SV nanoconjugates

ALS-raw, SV-raw, and ALS-SV nanoconjugate effects on PANC1 cell cycle analysis were investigated by flow cytometry. Cells were treated with different samples for 24 h. After that, cells were centrifuged and treated with 70% cold ethanol, washed, and then centrifuged. Propidium iodide and RNAse were mixed with the obtained separated cells before proceeding for flow cytometric analysis [21–23].

### Annexin V staining analysis

Comparative apoptotic potential of ALS-SV nanoconjugate was estimated, employing Annexin V staining. For performing this experiment, PANC1 cells were incubated in a 6-well plate (1×105 cells/well) and treated with ALS alone, SV alone, and ALS-SV nanoconjugate at the IC50 of different samples for 24 h at 7°C. Cells were then centrifuged at 200×g for 5 min., and the obtained cells were further suspended in PBS at room temperature. Now the cells were treated with ten µL of Annexin V and five µL of propidium iodide and left for incubation for five 25°C min. Finally, the obtained samples were visualized and analyzed using a flow cytometer (FACS Calibur, BD Bioscience, CA, USA) in triplicate [24,25].

### Caspase 3 estimation

A colorimetric assay kit (BioVision, Milpitas, CA, USA) was used for the estimation of caspase 3 level in treated PANC1 cells with various samples. For this purpose, PANC1 cells were cultured (3×106 cells/well) and treated with various samples such as normal saline, ALS alone, SV alone, and ALS-SV nanoconjugate. In addition, lysate buffer (ice-chilled) was used to suspend treated cells and left for incubation in ice for ten minutes. After ten minutes, cells were centrifuged at 10,000×g for 1 min, and obtained samples were used for the caspase-3 estimation as per the manufacturer’s instructions, and a microplate reader analyzed the evolved color at 405 nm [23,26,27].

### Real-time polymerase chain reaction (RT-PCR) for estimation of Bcl-2, Bax, *p53*, caspase 3 and TNF-α

The PANC1 cells were treated with ALS alone, SV alone, and ALS-SV nanoconjugate. The cell fraction was used for the extraction of RNA and proceeded for the synthesis of cDNA. Primer for the Bcl-2, Bax, p53, caspase 3, and TNF-α was designed by using Gene Runner software (Table 2) and the samples were normalized with β actin [11,22].

**Table 2.**
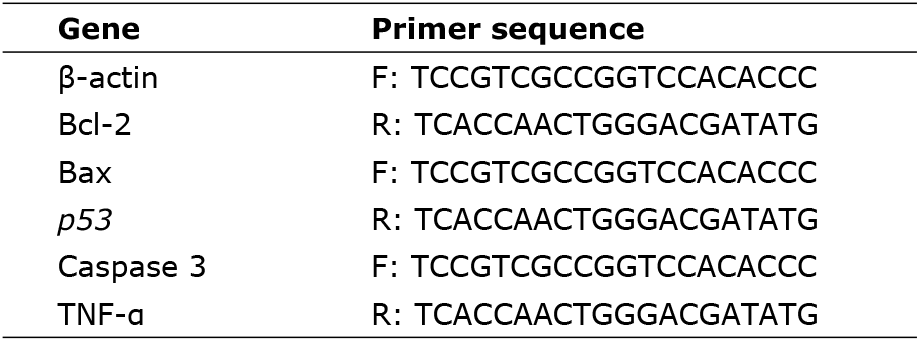
Nucleotide sequences for the primers used in the mRNA expression analysis.

### Determination of mitochondrial membrane potential (MMP)

MMP was investigated by ABCAM assay kit (Cambridge, UK) and PANC1 cells 5×103 (96 well plate) was incubated for 24 h. ALS alone, SV alone and ALS-SV nanoconjugate were separately added. The resultant cell mixture was kept in the dark, probe solution (tetramethylrhodamine, methyl ester) was replaced, and MMP was observed using FACS Caliber, BD Bioscience flow cytometer [28,29].

### Statistical analysis

Mean ± standard deviation (SD) was the expressed values. One Way ANOVA (followed by Tukey’s multiple comparisons test) was utilized for statistical analysis (p-value < 0.05).

## Results

### Experimental design-based optimization and selection of ALS-SV nanoconjugate

As per the runs (15 runs) obtained from the software, which was produced with the help of selected independent variables such as ALS: SV molar ratio, incubation time (min), and sonication time (min), various ALS-SV nanoconjugates were prepared (Table 3). Next, for the selection of optimized nanoconjugate, all the prepared ALS-SV nanoconjugates were characterized for particle size and zeta potential.

**Table 3.** Experimental matrix of the prepared formulations as suggested by the Box–Behnken design, with the values of the responses; particle size and zeta potential.

| Run | X<sub>1</sub>: ALS: SV molar ratio | X<sub>2</sub>: Incubation time (min) | X<sub>3</sub>: Sonication time (min) | Y<sub>1</sub>: Particle size (nm) | Y<sub>2</sub>: Zeta potential (mV) |
| --- | --- | --- | --- | --- | --- |
| 1 | 1: 8 | 12.5 | 0.75 | 76.8±2.1 | 12.3±0.2 |
| 2 | 1: 15 | 20 | 0.75 | 123.5±7.5 | 3.6±0.98 |
| 3 | 1: 8 | 5 | 0.5 | 76.3±7.6 | 11.3±0.9 |
| 4 | 1: 8 | 12.5 | 0.75 | 77.1±2.1 | 12±1.1 |
| 5 | 1: 8 | 5 | 1 | 67.8±3.2 | 13.6±0.4 |
| 6 | 1: 8 | 20 | 1 | 89±7.6 | 12.9±1.1 |
| 7 | 1: 1 | 12.5 | 0.5 | 49.6±3.4 | 21.7±2.1 |
| 8 | 1: 15 | 12.5 | 0.5 | 135±7.4 | 2.3±0.5 |
| 9 | 1: 1 | 12.5 | 1 | 42.4±8.6 | 23.2±2.3 |
| 10 | 1: 8 | 12.5 | 0.75 | 79.3±9.8 | 13.1±2.3 |
| 11 | 1: 15 | 5 | 0.75 | 110.5±11.1 | 5.4±1.1 |
| 12 | 1: 1 | 20 | 0.75 | 59.5±5.2 | 22.1±4.3 |
| 13 | 1: 1 | 5 | 0.75 | 37.8±2.3 | 26.6±2.2 |
| 14 | 1: 15 | 12.5 | 1 | 107.4±9.4 | 5.9±1.3 |
| 15 | 1: 8 | 20 | 0.5 | 101.2±5.5 | 10.6±1.1 |

### Impacts of independent variables on particle size

The statistical analysis of variance of response data (particle size) has been presented in Table 4. The software produced p-value declared the statistically significant effects of independent variables on selected responses of ALS-SV nanoconjugates. The independent factor (X1: ALS: SV molar ratio) exhibited a positive impact on particle size. Whereas independent factors X2 and X3 showed a comparatively shorter effect than X1 on particle size. Additionally, the R2 value and the value of R2 adjusted for particle size were found to be 99.4201 and 98.3764, respectively.

**Table 4.** Statistical analysis of variance (ANOVA) of the responses (Y_1_ and Y_2_) results.

| Factor<br>s | Y<sub>1</sub>: Particle size (nm) |  |  | Y<sub>2</sub>: Zeta potential (mV) |  |  |
| --- | --- | --- | --- | --- | --- | --- |
|  | Estimate | F ratio | P value | Estimate | F ratio | P value |
| **A:X<sub>1</sub>** | 6.995 | 754.44 | 0.0000 | -2.262 | 754.44 | 0.0000 |
| **B:X<sub>2</sub>** | 0.922 | 59.76 | 0.0006 | -0.405 | 59.76 | 0.0006 |
| **C:X<sub>3</sub>** | -77.669 | 28.19 | 0.0032 | 20.65 | 28.19 | 0.0032 |
| **AA** | 0.052 | 1.77 | 0.2409 | 0.032 | 1.77 | 0.2409 |
| **AB** | -0.041 | 1.39 | 0.2921 | 0.013 | 1.39 | 0.2921 |
| **AC** | -2.914 | 7.62 | 0.0398 | 0.3 | 7.62 | 0.0398 |
| **BB** | 0.045 | 1.74 | 0.2449 | 0.007 | 1.74 | 0.2449 |
| **BC** | -0.493 | 0.25 | 0.6379 | 0.0 | 0.25 | 0.6379 |
| **CC** | 52.933 | 2.96 | 0.1460 | -<br>12.133 | 2.96 | 0.1460 |
| **R<sup>2</sup>** | 99.4201 |  |  | 99.3752 |  |  |
| **Adjusted R<sup>2</sup>** | 98.3764 |  |  | 98.2505 |  |  |

At the end of software-based statistical analysis of particle size data, a polynomial equation was found (equation 1). The effects of multiple independent factors on particle size (dependent factor) can be recognized by the software-produced polynomial equation. In the equation, the positive sign recommended the positive effects of factors on the response. Exactly the same was seen in this study, where the particle size of nanoconjugates was increased with the increased molar ratio of ALS and SV (+6.99498) and incubation time (+0.922169). Meanwhile, the equation demonstrated the negative effect of sonication time (−77.669) on the particle size, where the particle size decreased with the increase of sonication time.

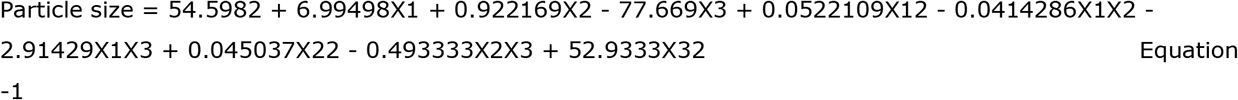

Preto chart (Figure 1.a) of particle size analysis showed a significant impact of independent variables and their interconnected impacts on the particle size of nanoconjugates. In Figure 1.a Preto chart distinctly demonstrated the positive effect of factors X1 and X2, and the negative effect of factor X3 over particle size. At the same time, software yielded contour plot (Figure 2) affirmed the Preto chart, which stated remarkable impacts of independent variables over particle size of nanoconjugates.

**Figure 1.**
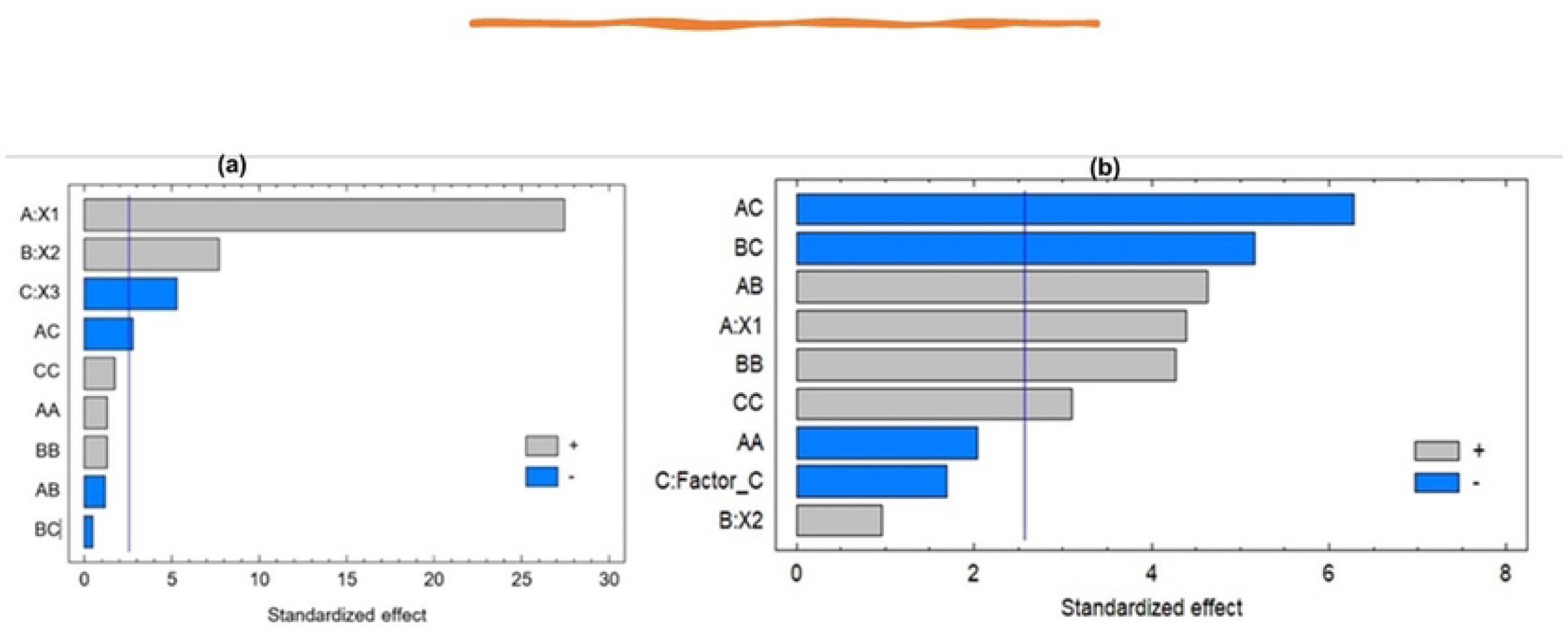
Preto chart for (a) particle size and (b) zeta potential, where X_1_, X_2_, and X_3_ represent the ALS: SV molar ratio, incubation time (min), and sonication time (min).

**Figure 2.**
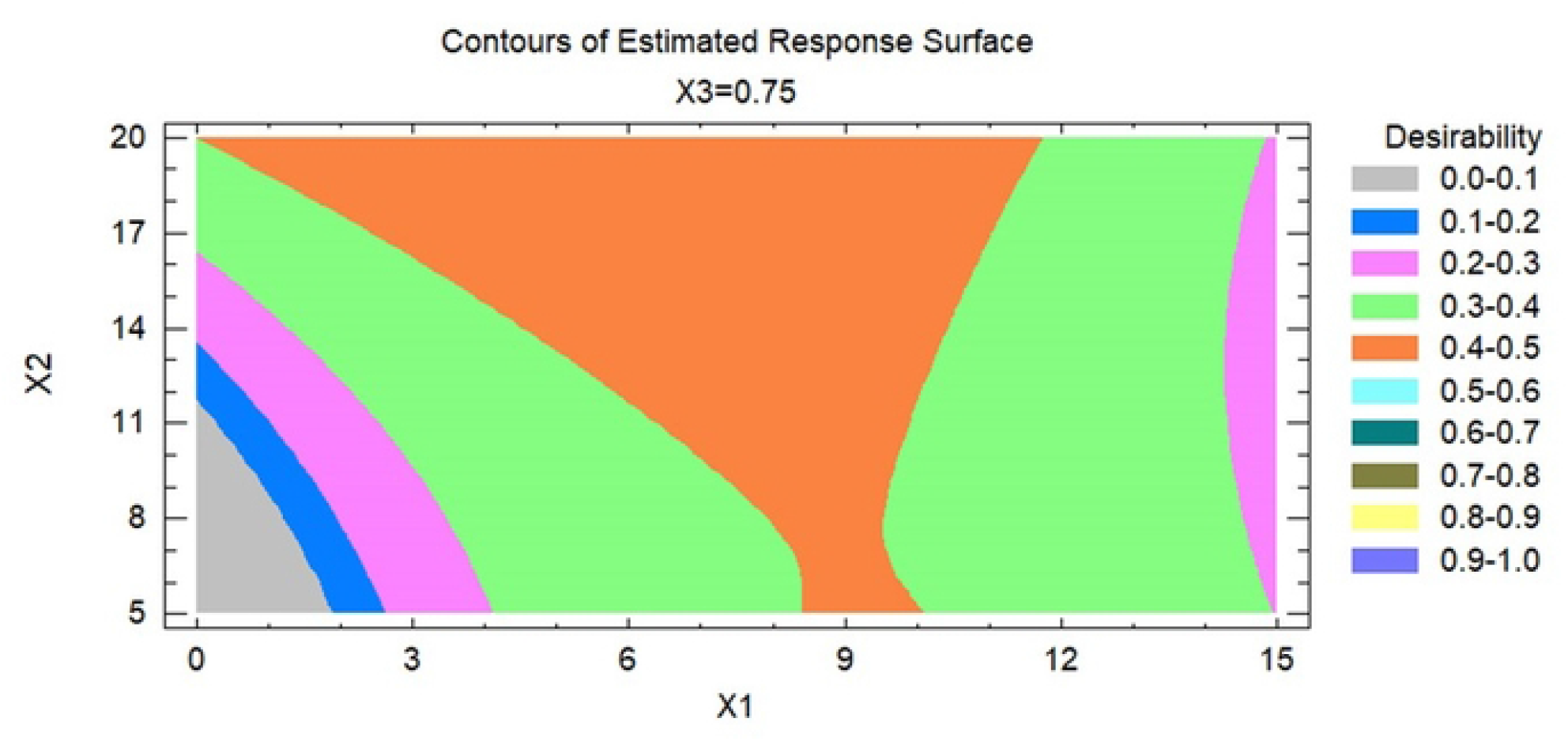
Contour plot of estimated responses, i.e., particle size (Y1) and zeta potential (Y2).

### Impacts of independent variables on zeta potential

The statistical analysis of variance of response data, i.e., zeta potential, has been presented in Table 2. The software produced p-values exhibited the statistically significant effects of independent variables on the zeta potential of ALS-SV nanoconjugates. The independent factor (X1: ALS: SV molar ratio) exhibited a negative impact on particle size and zeta potential. However, simultaneously independent factors X2 and X3 showed comparatively less effect than X1 on zeta potential. Additionally, the R2 value and the values of R2 adjusted for zeta potential were found to be 99.3752and 98.2505, respectively.

After software-based statistical analysis of zeta potential data, a polynomial equation was found (equation 2). The effects of multiple independent factors on zeta potential (dependent factor) can be confirmed by the software-produced polynomial equation. In the equation, the negative sign indicated the negative impacts of independent factors on the response. Exactly the same phenomena were seen in this study, where the zeta potential of nanoconjugates was increased with the decreased molar ratio of ALS and SV (−2.26156) and incubation time (−0.405265). Meanwhile, the equation demonstrated the positive effect of sonication time +20.65) on the zeta potential, where the zeta potential increased with the increase of sonication time.

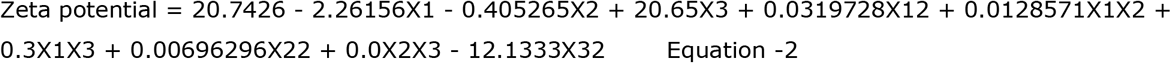

Preto chart (Figure 1.b) of zeta potential analysis showed a significant impact of independent variables and their interconnected impacts on the zeta potential of nanoconjugates. For example, in Figure 1.b Preto chart distinctly demonstrated the negative effect of factors X1 and X2 and the positive effect of factor X3 over zeta potential. At the same time, software yielded contour plot (Figure 2) affirmed the Preto chart of zeta potential, which declared remarkable impacts of independent variables over zeta potential of nanoconjugates.

### Optimization of ALS-SV nanoconjugate

After implementing the Box-Behnken response surface design, optimized ALS-SV nanoconjugate was selected with the minimum particle size and maximum zeta potential (Table 5).

**Table 5.**
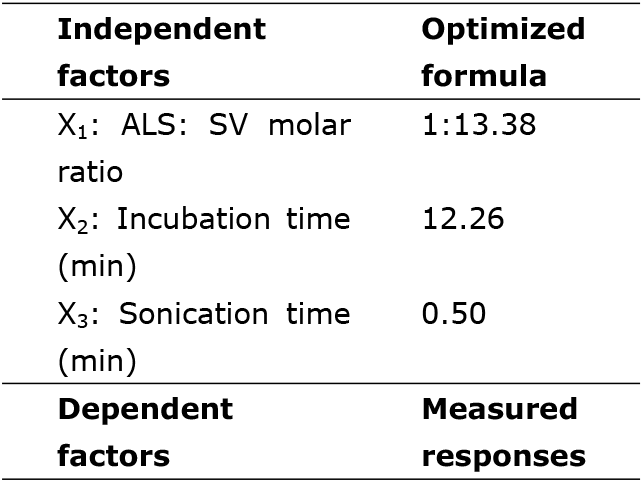
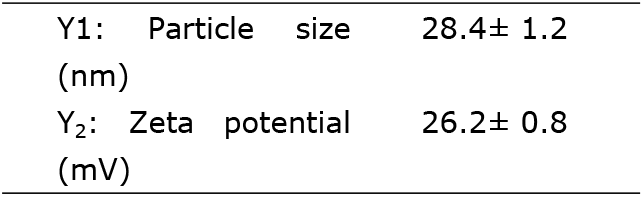
Optimized formula for ALS: SV nanoconjugate and its predicted responses.

### Cytotoxicity study of optimized ALS-SV nanoconjugate for determination of IC_50_

A comparative IC50 analysis was carried out via cell viability study through MTT assay on the PANC1 cell line, in which ALS alone, SV alone, and optimized ALS-SV nanoconjugate were taken as samples (Figure 3). The IC50 of ALS alone, SV alone, and optimized ALS-SV nanoconjugate was 22.3 ± 0.2 µg/mL, 6.23 ± 0.4 µg/mL, and 2.73 ± 0.2 µg/mL, respectively. Therefore, it can be concluded that theIC50 of ALS-SV nanoconjugate was significantly (p < 0.05) better than alone ALS and SV.

**Figure 3.**
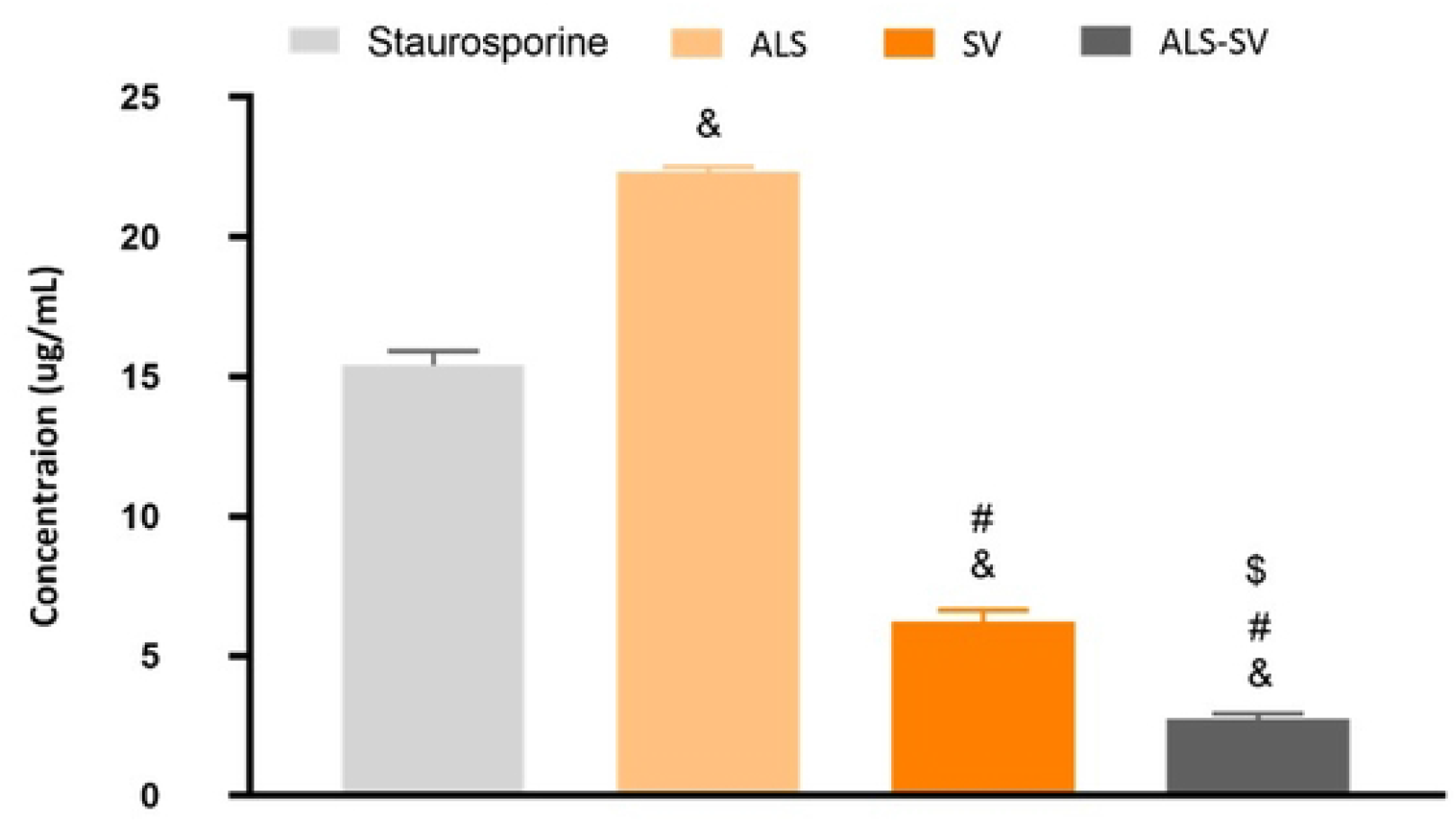
Results of cytotoxicity study of ALS alone, SV alone, and nanoconjugate of ALS-SV over PANC1 cell line. From these results, IC50 of samples was calculated after 24 h treatment by MTT assay. Data are the mean of 4 independent experiments ± SD. $ Significantly different vs SV, p < 0.05; # Significantly different vs ALS, p < 0.05; & Significantly different vs Staurosporine, p < 0.05.

### Determination effects of ALS-SV nanoconjugate on cell cycle

PI is a commonly used dye that binds with the DNA and is hence used for the estimation of the content of DNA during cell division. In the current study use of SV alone showed enhanced cell percent of the cell (53.18) arrested in the percent G0-G1 phase followed by the ALS-SV nanoconjugate (49.58%) and ALS alone (42.13%). However, Figure 4 showed a reduced percentage of the cell (29.15%) entering into the S phase after ALS alone treatment, followed by SV alone (36.91%) and the ALS-SV nanoconjugate (42.16%). However, ALS-SV nanoconjugate lowers the percent of cells entering the percent G2/M phase (8.26%). Followed by SV alone (9.91%) and ALS alone (28.72%), and hence, it can be concluded that ALS-SV was most active as an anticancer agent in the G2-M phase as presented in table 6.

**Table 6.** Results of the effect of samples on cell cycle phases.

| S. No. | Sample data code | IC50 (ug/mL) | DNA content in different phases (%) |  |  |  | Comment |
| --- | --- | --- | --- | --- | --- | --- | --- |
|  |  |  | G0-G1 | S | G2/M | Pre-G1 |  |
| 1 | ALS/PANC1 | 22.3±0.2 | 42.13±1.8 | 29.15±1.3 | 28.72±1.6 | 21.9±1.1 | Cell growth arrest@G2/M |
| 2 | SV/PANC1 | 6.289±0.4 | 53.18±1.4 | 36.91±1.3 | 9.91±2.1 | 26.08±1.1 | Cell growth arrest@ S |
| 3 | ALS-SV/PANC1 | 2.767±0.2 | 49.58±2.3 | 42.16±1.01 | 8.26±1.9 | 42.11±1.6 | Cell growth arrest@ S |
| 4 | Control PANC1 | -- | 51.03±2.5 | 44.13±1.1 | 4.84±2 | 2.03±1.01 | -- |

**Table 7.**
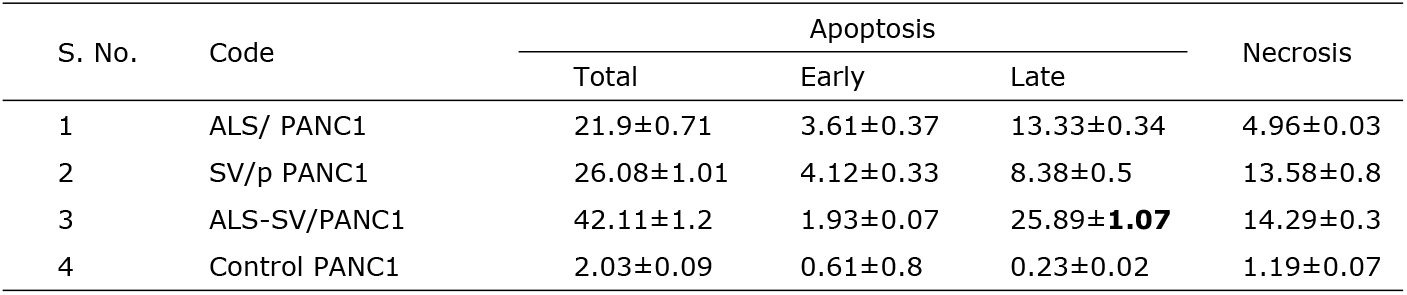
Results of apoptosis analysis via determination of Annexin V staining.

| S. No. | Code | Apoptosis |  |  | Necrosis |
| --- | --- | --- | --- | --- | --- |
|  |  | Total | Early | Late |  |
| 1 | ALS/ PANC1 | 21.9±0.71 | 3.61±0.37 | 13.33±0.34 | 4.96±0.03 |
| 2 | SV/p PANC1 | 26.08±1.01 | 4.12±0.33 | 8.38±0.5 | 13.58±0.8 |
| 3 | ALS-SV/PANC1 | 42.11±1.2 | 1.93±0.07 | 25.89±**1.07** | 14.29±0.3 |
| 4 | Control PANC1 | 2.03±0.09 | 0.61±0.8 | 0.23±0.02 | 1.19±0.07 |

**Figure 4.**
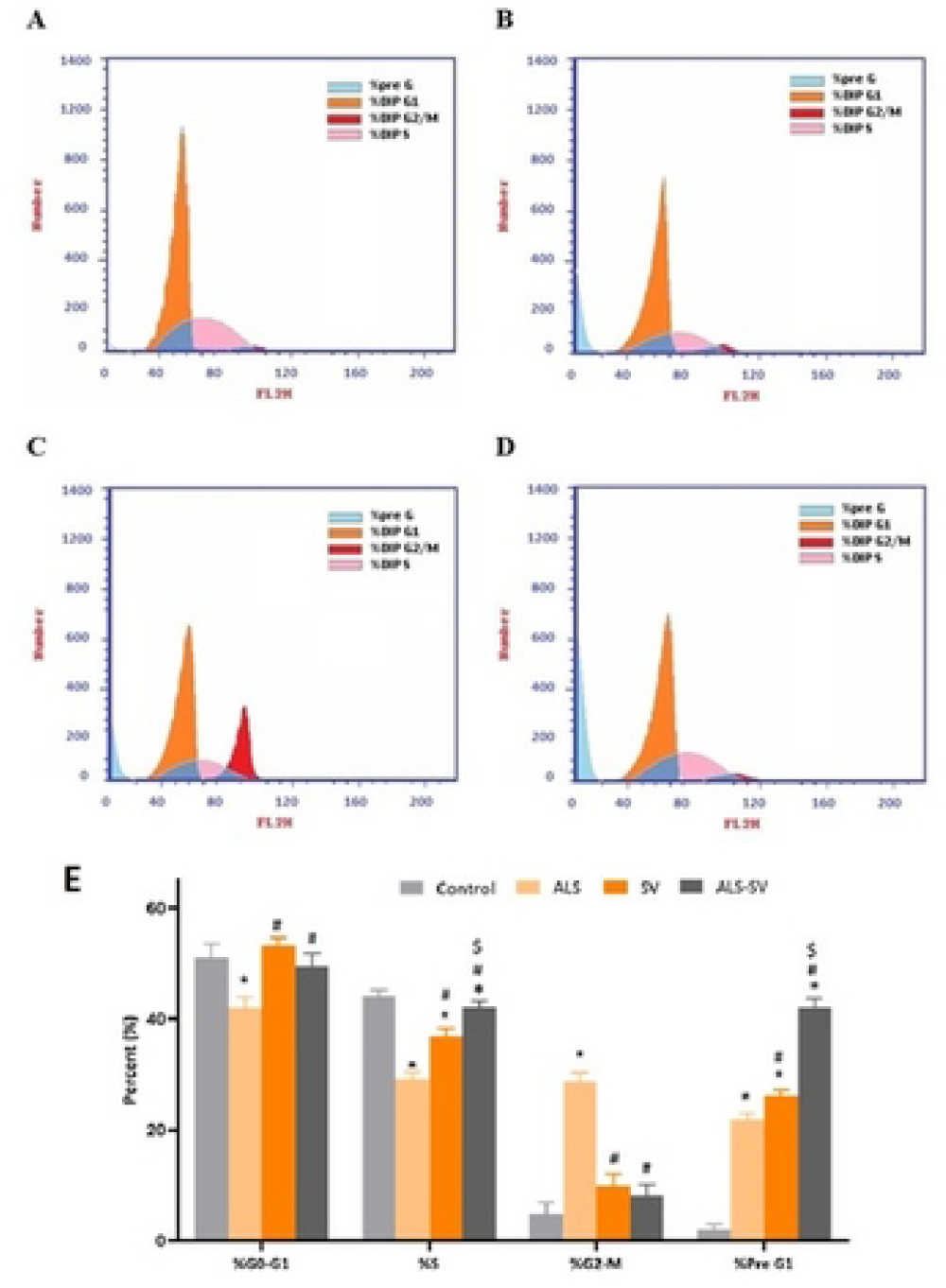
The flow cytometry analysis of cell cycle after PANC1 cells treated with ALS alone, SV alone, and ALS-SV nanoconjugate. *Significant vs control, p < 0.05; # significant vs ALS, p < 0.05; $ significant vs SV, p < 0.05.

### Analysis of apoptotic activity via Annexin V analysis

Apoptosis is one of the decisive parameters for the evaluation of the anticancer potential of the drug. Annexin V is commonly used in the assessment of apoptotic activity. This is because Annexin has a great affinity for the phosphatidylserine ligands located on the surface of cells and hence helps evaluate or identify the cells undergoing apoptosis. In the present study, ALS-SV nanoconjugate induced significant necrosis in the PANC1 cells, followed by ALS alone and SV alone. Additionally, ALS-SV nanoconjugate showed enhanced late apoptosis followed by SV alone and ALS alone. Moreover, ALS-SV also exhibited superior total apoptotic activity followed by ALS alone and SV alone. However, ALS alone showed the highest apoptotic activity in terms of early apoptosis, followed by SV alone and ALS-SV nanoconjugate (Figure 5). Thus, the study outcome showed superior apoptotic activity of ALS-SV during the late phase, and ALS showed early apoptotic activity,as in table 7.

**Figure 5.**
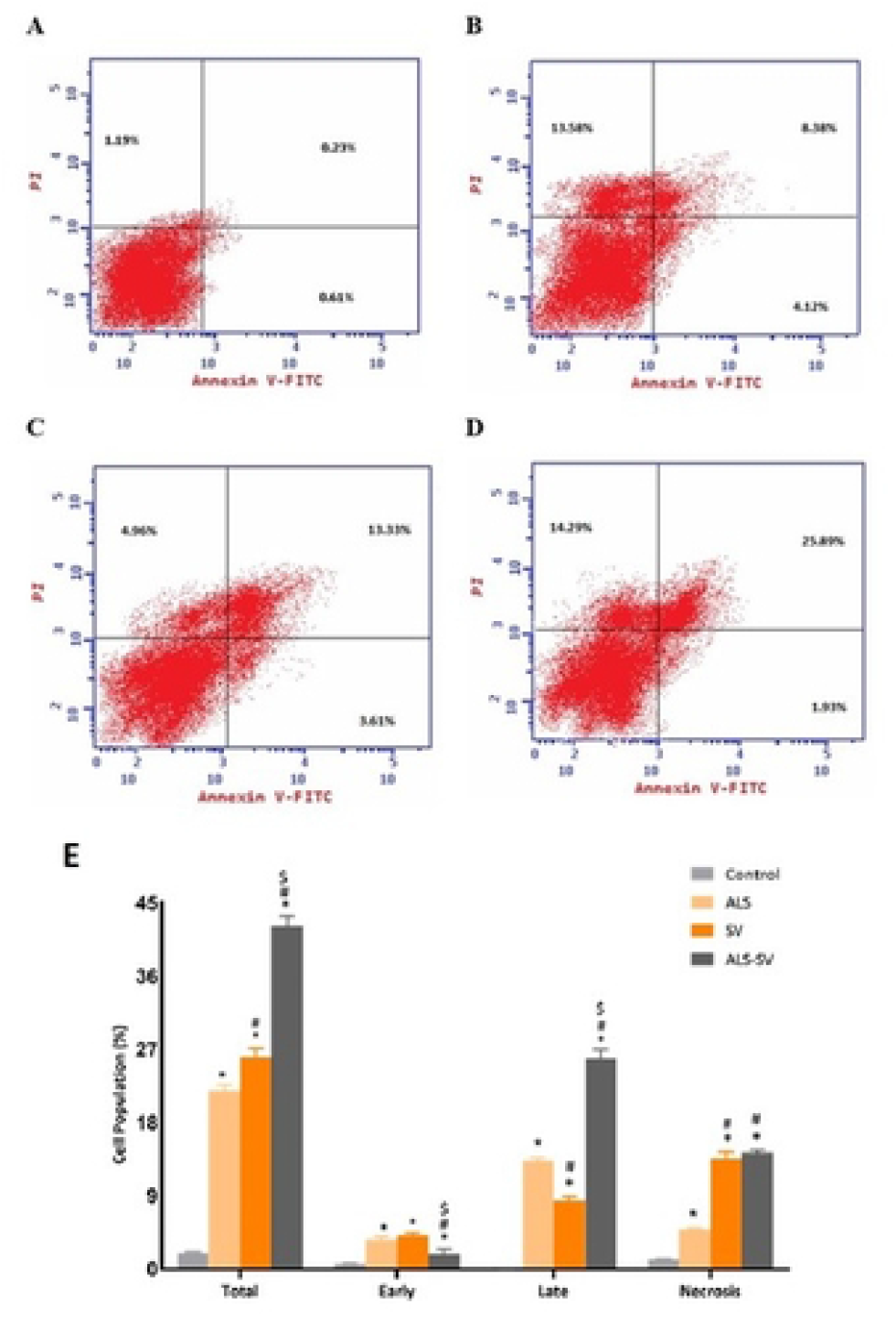
The flow cytometry analysis of apoptotic activity via Annexin V staining after PANC1 cells treated with ALS alone, SV alone, and ALS-SV nanoconjugate. ^*^Significant vs control, p < 0.05; ^#^ significant vs ALS, p < 0.05; ^$^ significant vs SV, p < 0.05.

### Estimation of Bax, Bcl-2, p53, and caspase 3

Apoptosis is one of the decisive parameters for anticancer activity. Therefore, a balance between pro and anti-apoptotic protein determines and anticancer potential of the drug. In the current study, the administration of ALS-SV nanoconjugate significantly increased the expression level of pro-apoptotic proteins such as Bax, caspase 3, and p53 and reduced the expression of anti-apoptotic protein Bcl-2 to ALS and SV alone (Figure 6). However, as compared to ALS alone, SV alone showed increased apoptotic activity.

**Figure 6.**
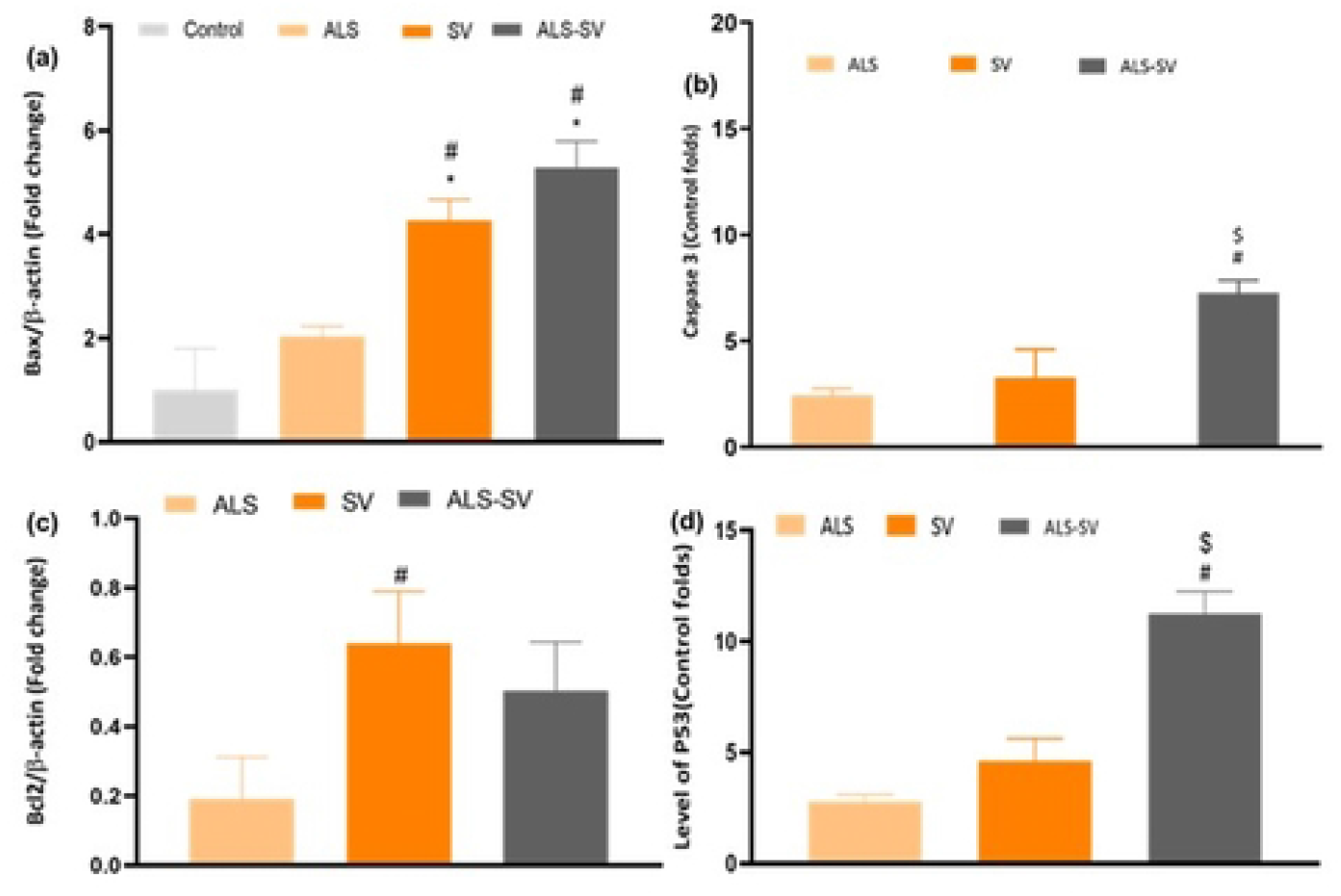
Showing the expression level of different pro and anti-apoptotic proteins such as (a) Bax, (b) caspase 3, (c) BCL-2, and (d) p53 against PANC1 cells when treated with ALS, SV alone and ALS-SV nanoconjugate. *Significantly different vs control, p < 0.05; # significantly different vs ALS, p < 0.05; $ significantly different vs SV, p < 0.05.

### Estimation of TNF-α

TNF-α is a well-known pro-inflammatory cytokine, and its increased level signifies cytotoxicity and anticancer activity [30]. In the current study, exposure to ALS-SV nanoconjugate treated cells demonstrated a significantly increased expression level of TNF-α as compared to ALS and SV alone (Figure 7).

**Figure 7.**
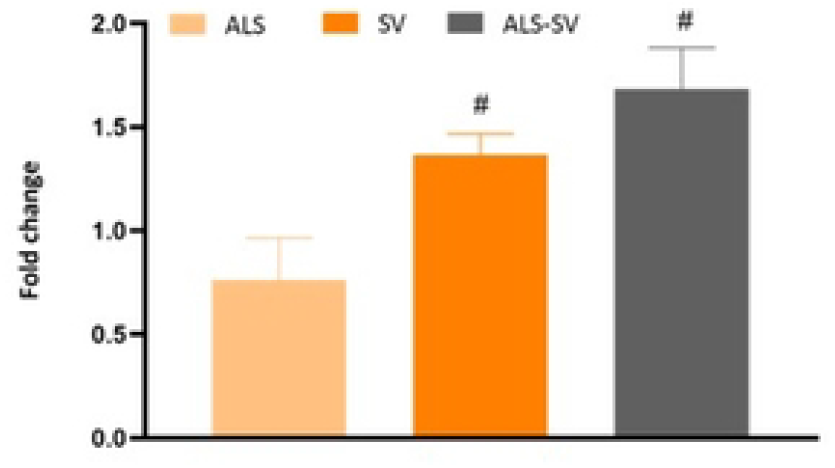
Showing the effect of ALS alone, SV alone, and ALS-SV nanoconjugate on the expression level of TNF-α when treated with the PANC1 cells. ^#^ significantly different vs ALS, p < 0.05.

### Determination of MMP

Change in the MMP occurred when the cell was exposed to the pro-apoptotic agents. Thus, the reduced mitochondrial potential is related to increased apoptosis. Currently, when PANC1 cells were treated with ALS-SV nanoconjugate, a lowered mitochondrial membrane permeability confirmed its apoptosis as well as anticancer effect (Figure 8).

**Figure 8.**
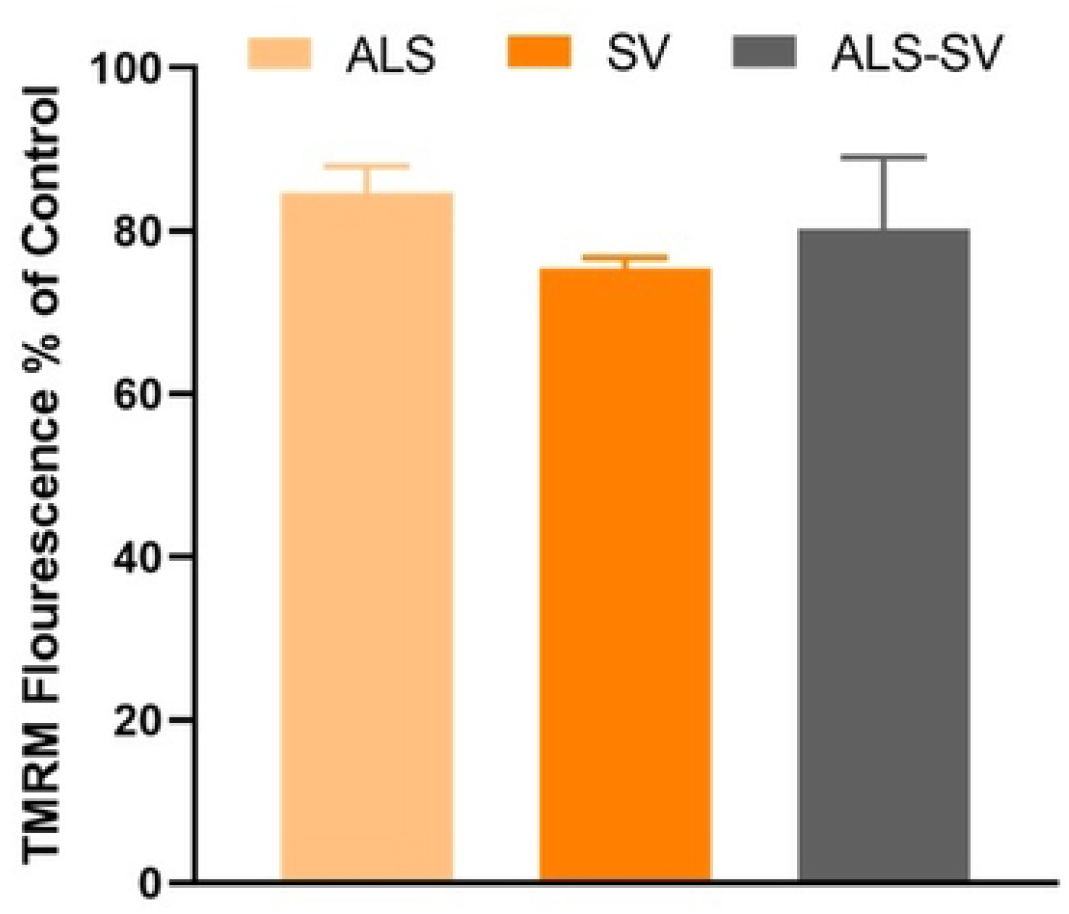
Showing the effect of ALS alone, SV alone, and ALS-SV nanoconjugate on the MMP when treated with the PANC1 cells.

## Discussion

In recent times, prevalence, as well as the incidence of PC, has increased exponentially. The concern with PC is the presence of asymptotic manifestation that often delays diagnosis that leads to tumor cells may reach the stage of metastasis. Also, the lack of therapeutic agents [2][4]. In addition, the deficiency of blood supply in the area of pancreas represent a hurdle against the bioavailability of chemotherapeutic agents in this area [4]. Thus, in the current study, nanoconjugate of ALS and SV was fabricated, optimized, and characterized. The anticancer potential of the formulation was validated in the PANC1 cells by performing various apoptotic parameters.

The optimized ALS-SV nanoconjugate formula was generated from the Box-Behnken experimental design. In this design, ANOVA-based statistical analysis yielded a positive significant (p < 0.05) impact of independent factors X1 (ALS-SV molar ratio concentration) and X2 (incubation time) on particle size as shown by the positive coefficient of equation 1. It means increased ALS-SV molar ratio and increased incubation time produced large size molecules due to enhanced conjugation. Whereas independent factors X3, i.e., sonication time, demonstrated a negative impact on particle size, as shown in the Preto chart (Figure 1.a). It means increased ultrasonication reduced the particle size of nanoconjugates which supports the previous statement about the effect of sonication time [31]. Simultaneously, the selected independent factors also affect the zeta potential of prepared nanoconjugates. For example, as shown in equation 2, the zeta potential of nanoconjugates was decreased with increment of ALS-SV molar ratio and incubation time, but at the same time, with the increment of sonication time, the zeta potential was also increased. Therefore, it can be concluded that the lower ALS-SV molar ratio and incubation time with increased sonication time produced the desired quality of nanoconjugate, which contained minimal size with maximized zeta potential that was required for better stability of prepared ALS-SV nanoconjugates.

The imbalance between the cell cycle and death and changes in the signaling pathways are what cause cancer [32]. During the 1970s, the first-time association between carcinogenesis and apoptosis was established [33]. In cancer, reduced apoptosis by pro-apoptotic gene downregulation and anti-apoptotic gene upregulation of have been reported [34]. Additionally, tumor cells develop the mechanism of apoptosis evasion, reduced activity of pro-apoptotic proteins, reduced caspases activity, and alteration in the functioning of death receptors [11,35].

The apoptotic proteins Bcl-2 and Bcl-xL have been linked to the survival of tumour cells by preventing the mechanism of programmed cell death. Furthermore, it was observed that Bcl2 overexpression was linked to proliferation and Myc-induced angiogenesis. [32,33]. The pro-apoptotic protein Bax, on the other hand, is connected to the induction of apoptosis and works in conjunction with caspases to cause the death of tumour cells. Therefore, an increase in Bcl-2 is linked to decreased oncogenic activity, whereas an increase in Bax is linked to enhanced oncogenic activity. p53 is a tumor suppressor gene and extensively explored in apoptosis-related to carcinogenesis. p53 is considered as a checkpoint protein involved in the arrest of the cell cycle via stimulation of DNA damage [33]. This work investigated the anticancer potential of Als alone, SV alone, and ALS-SV nanoconjugate. The results proved the enhanced pro-apoptotic and anticancer potential of ALS-SV nanoconjugate by showing decreased expression of Bcl-2 and increased expression of Bax and p53 in PANC1 cells.

It has been discovered that MMP and the permeability transition pore (PTP) responsible for the release of cytochrome C from the outer mitochondrial membrane are altered by pro-apoptotic proteins like Bax and pro-inflammatory cytokine TNF-α. [34,36].

When cytochrome C is distributed from the pore of mitochondria, it binds to Apaf-1 and initiates the cascade for the production of apoptosomes and caspases like caspase 3 then continue and cause apoptosis [36]. Furthermore, TNF-α binds to the receptors and starts the extrinsic apoptosis pathway that causes the activation of procaspase-8, which in turn activates caspase-8, which in turn converts procaspase-3 generated by Cyt c, Apaf-1, and apoptosome into caspase-3 and results in cell death [37,38]. In the current work, treatment of the PANC1 cells with optimized ALS-SV nanoconjugate resulted in decreased MMP, increased TNF-α and caspase-3 production, which indicates mitochondrial-mediated apoptosis. Furthermore, Annexin V assay confirmed apoptotic activity in the PANC1 cells and established the ALS-SV nanoconjugate’s late apoptotic and necrotic activity. It is generally known that an anticancer agent’s ability to arrest the cycle of cancer cells is confirmed by the therapeutic agent’s apoptotic potency. Utilization of optimized ALS-SV in the current investigation demonstrated cell cycle arrest at the G2-M phase and validated the anticancer potential of the nanocnjugate.

## Conclusions

This study produced an optimized ALS-SV nanoconjugate that met the criteria for choosing an optimum nanoconjugate using Box-Behnken-based design expert software, with the smallest size and highest zeta potential. Following that, the relative therapeutic efficacy of treating PANC1 cells with optimized ALS-SV nanoconjugate, ALS alone, and SV alone was determined. Then in order to determine the anti-PC effects, treated cells with various samples were analyzed for cytotoxicity, cell cycle arrest, annexin V, Bax, Bcl-2, p53, caspase 3, TNF-α, and MMP, which illustrated an excellent anti-PC activity by intercepting cell cycle in G2-M phase, and superior apoptosis. Thus, it was concluded that the optimized ALS-SV nanoconjugate had superior beneficial results, making it a novel therapeutic strategy for PC.

## Author Contributions

“Conceptualization, N.A. and M.W.; methodology, W.Y.; software, A.F.; validation, W.Y., A.F. and M.W.; formal analysis, W.Y.; investigation, W.Y.;; resources, N.A.; data curation, N.A..; writing—original draft preparation, W.Y.; writing—review and editing, W.Y.; visualization A.F.; supervision, N.A..; project administration, N.A.; funding acquisition, N.A.. All authors have read and agreed to the published version of the manuscript.

## Acknowledgments

In this section, you can acknowledge any support given which is not covered by the author contribution or funding sections. This may include administrative and technical support, or donations in kind (e.g., materials used for experiments).

